# Altered DNA methylation pattern contributes to differential epigenetic immune signaling in the upper respiratory airway of COVID-19 patients

**DOI:** 10.1101/2024.04.29.591494

**Authors:** Melissa Govender, Jyotirmoy Das, Francis R. Hopkins, Cecilia Svanberg, Johan Nordgren, Marie Hagbom, Jonas Klingström, Åsa Nilsdotter-Augustinsson, Yean K. Yong, Vijayakumar Velu, Sivadoss Raju, Johanna Sjöwall, Esaki M. Shankar, Sofia Nyström, Marie Larsson

## Abstract

The emergence of SARS-CoV-2 has had a profound adverse impact on global health and continues to remain a threat worldwide. The disease spectrum of COVID-19 ranges from asymptomatic to fatal clinical outcomes especially in the elderly population and in individuals with underlying medical conditions. The impact of COVID-19 on host immune responses and immune cells at the protein and DNA levels remains largely ambiguous. In a case-control study, here we explored the impact of COVID-19 on DNA methylation patterns in the upper respiratory airway to determine how SARS-CoV-2 infection altered the immune status of individuals requiring hospitalization for COVID-19. We performed DNA methylation arrays on nasopharyngeal swabs at inclusion/hospitalization as well as 6 weeks post-inclusion. Our study reveals a distinct DNA methylation pattern in COVID-19 patients compared to healthy controls, characterized by 317 779 differentially methylated CpGs. Notably, within the transcription start sites and gene body, COVID-19 patients exhibited a higher number of genes/CpGs with elevated methylation levels. Enrichment analysis of methylated genes at transcription start sites highlighted the impact on genes associated with inflammatory responses and immune functions. Some SARS-CoV-2 -induced CpG methylations were transient, returning to normal levels by 6 weeks post-inclusion. Enriched genes of interest included IL-17A, a pivotal cytokine implicated with inflammation and healing, and NUP93, associated with antiviral innate immunity. Further, six genes in our data set, OAS1, CXCR5, APP, CCL20, CNR2, and C3AR1, were found in enrichment analysis with previous COVID-19 studies. Additionally, RNAse1 and RNAse2 emerged as key regulators, while IL-18 played a role in various biological processes in COVID-19 patients. Overall, our results demonstrates that COVID-19 has a major impact on the upper airway by modifying the methylation pattern of many genes and this could have implications for the conditioning of the airways and how the individual response to future airway infections.

## Introduction

The rapid spread and consistent evolution of SARS-CoV-2 variants continues to widen the magnitude of health threat inflicted by COVID-19 on humankind. Since the onset of the COVID-19 pandemic in 2020, substantial advances have been achieved in the realm of therapeutics, together with the development of a wide array of vaccines to combat the SARS-CoV-2 pandemic ^1^. Despite increased control measures, the world has witnessed an high death toll due to SARS-CoV-2 infection largely among non-vaccinated individuals ^2^. Furthermore, the spectrum of symptoms and the long lasting negative effects among people with severe COVID-19 ^3, 4^ together with the viral evolution leading to possible new SARS-CoV-2 variants of concern ^5, 6^, continue to pose formidable challenges to effective disease prevention. The severity of COVID-19 has been linked to dysregulation of immune responses induced by an exaggerated and prolonged inflammation ^7^. Hence, a thorough understanding of the nature of immune responses elicited against SARS-CoV-2 is of paramount importance and relevance.

A large proportion of COVID-19 research has been devoted to phenotype, proteomic ^8, 9^ and/or transcriptomic studies ^10, 11, 12^, which has given vital information on the dynamics of the immune cell landscape heretofore. In a previous study, we revealed persistent alterations within the monocytic, dendritic cell, and T cell compartments induced by SARS-CoV-2 infection, which manifested even six to seven months post-hospitalization ^13, 14^. This underscores the enduring impact of the virus on immune cell dynamics, highlighting the necessity for continued exploration into the intricacies of the immune responses against SARS-CoV-2.

More recently, there have been multiple epigenetic studies aimed to unravel the complexities of SARS-CoV-2 pathogenesis and its enduring impact on the host ^15, 16, 17^. Of the various epigenetic modifications, DNA alterations, particularly methylation, exhibit remarkable stability, and form an integral component of a cell’s programming, persisting throughout its life cycle and divisions. The activation or silencing of certain genes by DNA modifications is one of the immune regulatory mechanisms implicated in the control of infection and attrition of inflammatory responses to avoid tissue damage in the host ^18^. The regulatory influence of DNA methylation extends to transcriptional accessibility of genes and the subsequent effects, i.e., activity and expression, depending on the region in the genome/gene affected ^19, 20^. Many biological and environmental factors appear to influence the DNA methylation patterns such as age, biological sex, and body mass index (BMI) ^21, 22^. Likewise, infections, both viral and bacterial, can elicit rapid epigenetic alterations, such as DNA methylation, and thereby regulate gene expression in cells ^5, 6, 23^.

Blood DNA methylation patterns in asymptomatic and mild COVID-19 cases differed from patterns found in healthy individuals, and genes such as Wnt, and signaling pathways such as muscarinic acetylcholine receptor signaling, and gonadotropin-releasing hormone receptor pathways were enriched in COVID-19 ^24^. Unique SARS-CoV-2-specific methylation patterns were found in COVID-19 patients compared to pre-pandemic healthy controls and to patients with other upper respiratory infections due to rhinovirus and influenza B virus ^25^. In addition, the immune functions in SARS-CoV-2-infected individuals are also affected by DNA methylation, and certain methylation profiles have been linked to disease severity, such as respiratory distress and intensive care unit (ICU) admission ^25, 26^.

COVID-19 patients with acute respiratory distress syndrome (ARDS) had 14% differently methylated genes in the promoter regions compared to healthy controls. These promoter regions regulate genes involved in regulating immune pathways, such as IFN-γ and IFN-α ^27^. In addition, higher methylation, i.e., hypermethylation of genes in the ‘apoptotic execution pathway’ has been linked to higher mortality risk in COVID-19 patients ^27^. Furthermore, several DNA methylations occurred in the inflammasome component absent in melanoma (AIM) gene during COVID-19 progression, which were linked to heightened immune responses among patients who recovered from SARS-CoV-2 infection ^27^.

Several epigenetic studies have investigated the methylation patterns in blood samples of individuals with mild to severe COVID-19 ^24^ ^25, 26^ ^27^. Given that SARS-CoV-2 infection is initiated in the upper airway, establishing a microenvironment involving the respiratory airway mucosa, and that immune cells of the airway mucosa considerably differ from those distributed in the blood, we aimed to explore the methylation status in the upper respiratory airway using nasopharyngeal specimens obtained from hospitalized patients with moderate to severe COVID-19. We observed a significant difference in the methylation patterns among COVID-19 patients at hospitalization/inclusion and at 6 weeks post-inclusion relative to healthy controls. We found that several genes were differentially methylated revealing significance at hospitalization as compared to 6 weeks post-inclusion. Unsupervised hierarchical clustering of DNA methylation β-values at both overall and transcription start sites revealed distinct differences between COVID-19 patients and controls. Enrichment analyses of methylated genes at transcription start sites in COVID-19 patients highlighted impacts on inflammatory response and immune processes. Some SARS-CoV-2 -induced CpG methylations were transient, reverting to normal levels within 6 weeks post-infection. At inclusion, certain unique gene ontology biological processes were observed, including response to stimulus and cellular process. Notably, the pro-inflammatory factor IL-17A was enriched at inclusion and the antiviral NUP93 factor was enriched at 6-week post inclusion. RNAse1 and RNAse2 emerged as top regulators as well as IL-18, a vital factor that influences various biological processes in COVID-19. This is, to our knowledge, the first instance of investigation of DNA methylation patterns in the nasal area in COVID-19 patients, providing new insight to immune events in the upper airways during COVID-19.

## Materials and methods

### Participants and sample collection

The study was approved by the Swedish Ethical Review Authority (Ethics No. 2020-02580) and was carried out in accordance with Good Clinical Laboratory Practices, the International Conference on Harmonization Guidelines, and the Declaration of Helsinki. The patient cohort included hospitalized COVID-19 patients (N=27; age range 26-91 years), and SARS-CoV-2-negative healthcare workers as healthy controls (HC) (N=12; age range 26-62 years), which were selected from recruited individuals previously described ^13, 14^. All individuals provided written informed consent prior to enrolment. The COVID-19 patient clinical data and additional information are described in **Table 1**.

**Table 1.**
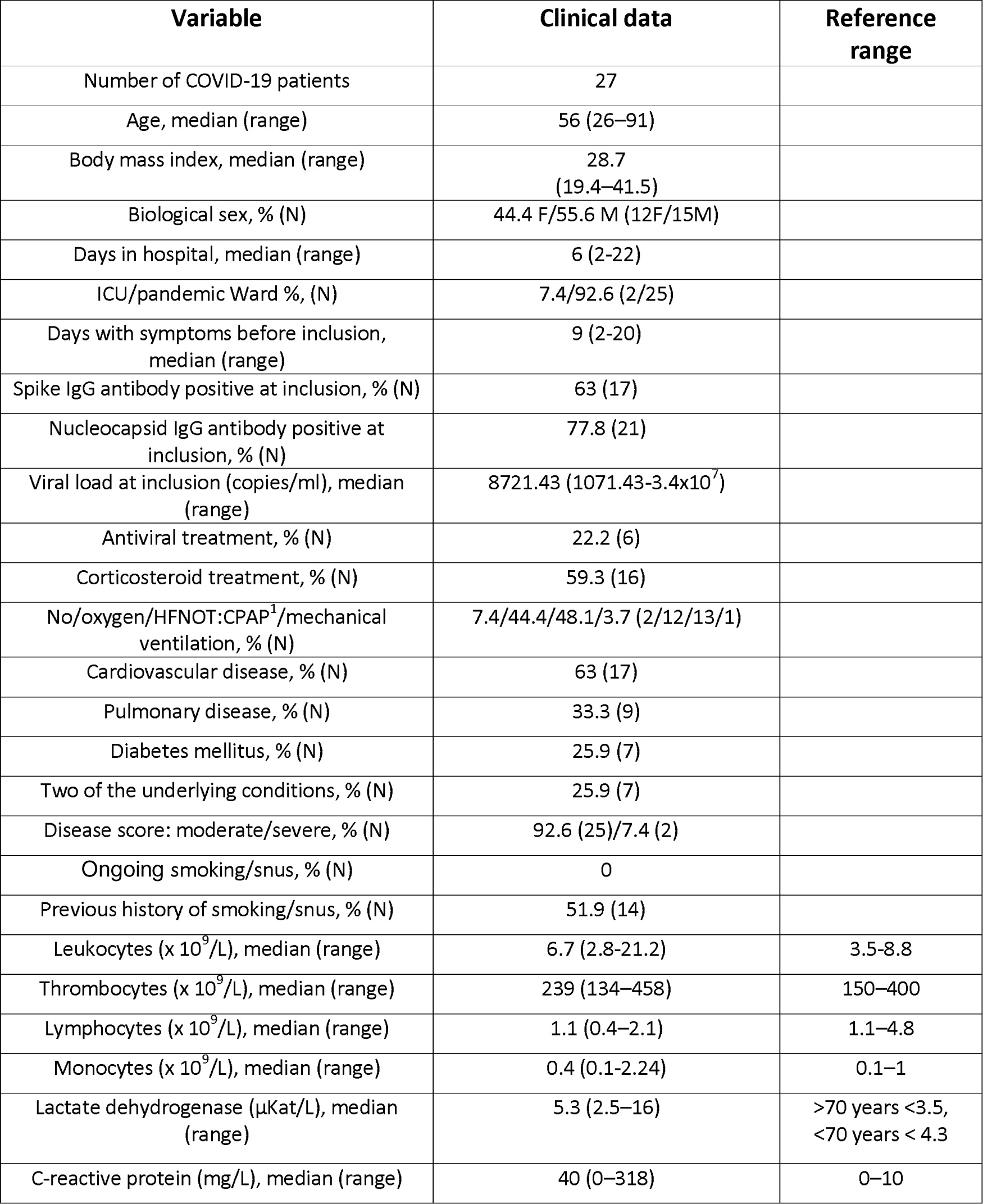
Clinical and-demographical characteristics of hospitalized COVID-19 patients.

COVID-19 disease score was defined according to the criteria advocated by the National Institutes of Health ^4, 28^, and slightly modified based on the requirement for supplementary oxygen and the highest level of care required. Patients in this cohort were classified mostly as moderate to severe based on the above criteria **(Table 1)**. Nasopharyngeal specimens were collected at inclusion (baseline) (timepoint 1) and at 6 weeks post-inclusion (timepoint 2). The samples were transferred into microfuge tubes and centrifuged at 1800 RPM for 6 minutes at 4°C, and the pellets were frozen at -80°C for the extraction of genomic DNA (gDNA) for later use in the experiments.

### Genomic DNA isolation

Genomic DNA (gDNA) was isolated using the Qiagen AllPrep DNA/RNA Mini kit (Qiagen) according to the manufacturer’s instructions. Briefly, 600 µl of lysis (RLY) buffer was added to thawed pellets and mixed. The lysate was transferred to the AllPrep DNA column and centrifuged at 11000 g for 1 min. The column was washed with 500 µl AW1 buffer (centrifuged at 11000 g for 1 min) and washed with 500 µl AW2 buffer. Finally, the columns were placed in 1.5 ml microfuge tubes, and 30 µl extraction buffer (EB) was added for a 1 min incubation, before eluting the gDNA by centrifuging at 11 000 g for 1 min. The elution step was repeated with the eluant to increase the gDNA concentration.

### Quantification of genomic DNA

The quality and concentration of the genomic DNA was measured using the broad range (BR) dsDNA Quantitation Qubit assay (Invitrogen, USA) according to the manufacturer’s instructions. Briefly, the samples were prepared with the Qubit working solution (Qubit reagent diluted 1:200 in Qubit buffer). The standards as well as nasopharyngeal samples were prepared in working solution and were vortexed for a few seconds before incubation for 2 min at room temperature. The amount of genomic DNA was then quantified by measuring the fluorescence with the Qubit™ dsDNA BR Assay Kit (Invitrogen, USA) on a Qubit^TM^ 2.0 Fluorometer (Invitrogen^TM^ by Life Technologies^TM^, Thermofisher, USA).

### Sample preparation for DNA methylation microarray

Quantified genomic DNA samples (69-552 ng) were transported to the Bioinformatics and Expression Analysis Core facility (BEA) based at the Karolinska Institute, Stockholm, Sweden. Here, the samples were subjected to bisulfite conversion using the Zymo Research EZ-96 DNA Methylation™ Kit (D5004). The samples were quantified with the Qubit assay, and 70-250ng was used as input for the Illumina Infinium Methylation EPIC 850K BeadChip array (Illumina Inc., San Diego, USA) following the protocol outlined by Illumina. QC, normalized data, and IDAT files were obtained and used for further analysis.

### Data processing and statistical analysis: Differential methylation analysis

The IDAT files from Illumina® HumanMethylation EPIC arrays were analyzed using R (v4.2.1)^32^ and Bioconductor packages (v3.16), Chip Analysis Methylation Pipeline (ChAMP)^33^ analysis package (v2.28.0). The data initially contained 865918 probes. The files were pre-processed to filter out CpGs with detection *p* value >0.01 (removing 32515 probes), with a bead count of <3 in at least 5% of samples (removing 5192 probes), all non CPG probes and only keeping CpGs (removing 5192 probes); those with SNPs ^30^ (probes that align to multiple locations as identified in Nordlund et al ^31^; and finally, filtering probes located on X and Y chromosomes. 715410 probes and all 48 samples were left remaining for analysis.

. A quality assessment on the filtered data was performed and using beta-mixture quantile normalization (BMIQ) function and normalized dataset was calculated. The β- and M-values for each CpG per sample was estimated in a distribution plot. To reduce the batch effect in relation to biological variation on the data matrix, deconvolution (singular value decomposition, SVD) was performed on the normalized data using *runCombat* function and corrected against the confounding factors lactate dehydrogenase (LDH), and BMI. The differential methylation analysis was done on the corrected data with the linear modeling (*lmFit*) and *eBayes* algorithm between two sample groups. The differentially methylated CpGs were considered significant at the Bonferroni-Hochberg (BH)-corrected *p* value (*p* value_BH_) <0.05. The hierarchical cluster analysis was performed using the Euclidean distance calculation within the *ape* package ^34^(v5.7). The principal component analysis was performed using *FactoMineR* ^35^(v2.8) and *factoExtra* ^36^ (v1.0.7) packages with *in house* R script.

### Downstream analysis: Feature analysis of differentially methylated CpGs

The resulted differentially methylated CpGs were annotated using *AnnotationDbi* package ^37^ (v1.60.2) (Human Genome version 38) using the *in house* script to visualize the genomic distribution. The volcano plot displayed the distribution of hypermethylated and hypomethylated (log_2_FC cut-off <0.3) significant differentially methylated CpGs (*p* value_BH_ <0.05) using *in house* R script with ggplot2 package ^38, 39^ (v3.4.2). The cut-off score was calculated using the β-value distribution of all samples with mean±2SD. Heatmaps were calculated using in-house R script with Complex. Heatmap package ^40^(v2.14.0) from individual β-value. The differentially methylated CPGs result was filtered based on the genomic location, selecting the transcription start site (TSS) regions (TSS200 and TSS1500) and gene body regions using log2FC cut-off score of >0.3 for hypermethylated and <0.3 for hypomethylated and genomic location. Separated heatmaps were created to visualize the β-value distribution. The differentially methylated CpGs were annotated with their respective official gene symbols.

Enrichment analysis was performed using the Metascape database (http://metascape.org){Zhou, 2019 #3} to determine any biological significance of the pooled 1000 differentially methylated genes, i.e. top 500 hypomethylated and 500 hypermethylated genes, in the transcription start site regions TSS1500 and TSS200 in the COVID-19 patients. Further analysis was performed using the CORONASCAPE database (https://metascape.org/COVID/){Zhou, 2019 #3} to identify any similarities between our data, i.e. the pooled 1000 differentially methylated genes, and differentially methylated genes from other COVID-19 datasets. The Ingenuity Pathway Analysis (IPA) software was used for pathway, top regulator analysis and network summary of the differently methylated genes in the TSS1500 and TSS200 regions. The data set included 5077 mapped ID that gave 2831 analysis-ready molecules with log FC -0.5503 to 0.5475.

### Statistical analysis

All differences with a *p* value_BH_ <0.05 were considered significant, if not stated otherwise. We calculated family-wise error rate (FWER) using the Bonferroni-Hochberg (BH)-correction method. All analyses were performed using R (v4.2.1) with the aforementioned packages.

## Results

### Patients and clinical characteristics

To assess the DNA methylation profiles in the upper respiratory airways of hospitalized COVID-19 patients (N=27), a total of 36 nasopharyngeal swabs were collected, with 21 samples from hospitalization/inclusion (T1), and 15 samples 6 weeks post-inclusion (T2). In addition, nasopharyngeal swabs from healthy subjects (N=12) were used as controls **(Figure 1)**. Participants were recruited between July 2020 to October 2021, and represented hospitalized patients with moderate to severe COVID-19 manifestations, as determined by the Guidelines of the National Institutes of Health, based on the maximum oxygen required and the highest level of care needed ^28^. At this time the following SARS-CoV-2 strains circulated in Sweden: alpha, beta, gamma, and delta. Of the total patients, 25.9% had at least two of the following underlying conditions, i.e., cardiovascular disease, pulmonary disease, and diabetes mellitus. The median number of days with symptoms prior to inclusion into the study was nine **(Table 1)**. Nasopharyngeal swabs (N=12) were obtained from healthy controls at enrollment. Standard clinical and blood parameters were assessed at inclusion for COVID-19 patients **(Table 1)** as well as controls.

**Figure 1:**
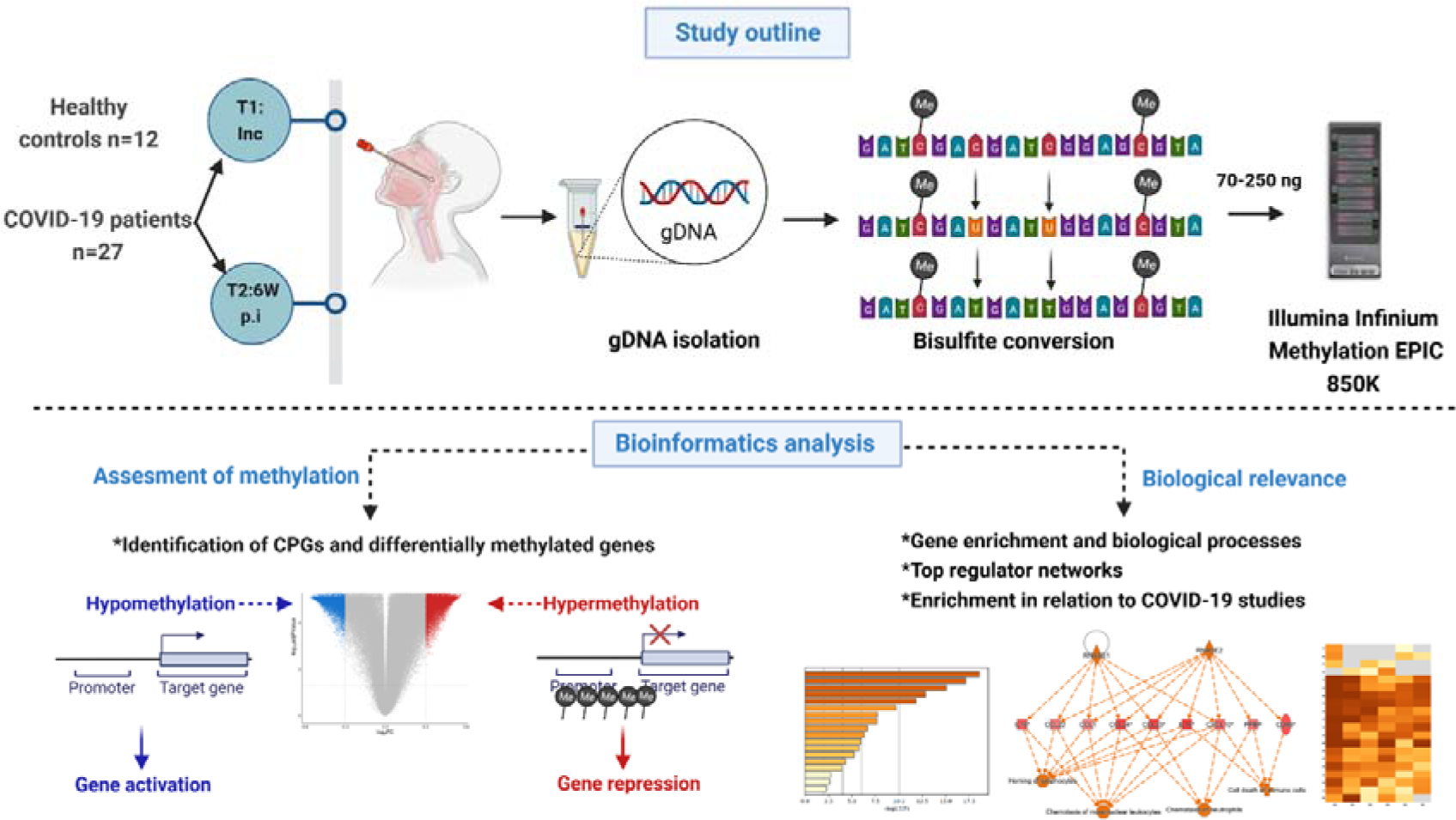
Graphical summary of the study and methods. Nasopharyngeal samples were collected from 27 COVID-19 patients at the beginning of the study, i.e., inclusion, time point 1(T1); and followed up 6 weeks later (T2, 6W post inclusion), (N=36). Healthy controls (HC) (N=12) were included at inclusion. Genomic DNA (gDNA) was isolated and sent for bisulfite treatment and methylation analysis with the Illumina infinium Methylation EPICS 850K. Bioinformatics analysis was performed to identify unique CPGs and differentially methylation genes, followed by further analysis to assess the biological relevance.

### Raw DNA methylation data revealed high quality and beta distribution

Genomic DNA (gDNA) from nasopharynx samples were prepared for bisulfite conversion, and we performed a DNA methylation array (Illumina Infinium Methylation EPIC 850K). The data were processed and filtered with bioinformatics analysis to identify methylation patterns and determine their biological significance **(Figure 1)**. Initial quality control assessment of the raw methylation data indicated a good continuous probability distribution, i.e., beta distribution, between samples **(Supplementary Figure 1A)**, and separation of healthy control samples and inclusion and 6-week time-points samples from COVID-19 patients in the multidimensional scaling plot **(Supplementary Figure 1B)**. Lactate dehydrogenase and BMI were estimated as significant components of variation/confounding factors in the methylation dataset, and the effect of these factors were corrected using singular value decomposition, i.e., deconvolution. LDH was disregarded due to being a patient-related factor, and the dataset was adjusted for the BMI **(Supplementary Figure 1C)**.

### COVID-19 patient DNA methylation pattern differs from that of healthy controls

For the identification of differentially methylated CpG sites, an assessment of the samples from COVID-19 patients, both at inclusion and 6-week timepoints, and healthy controls samples was performed. A total of 317779 statistically significant differentially methylated CpGs were identified in the pooled COVID-19 patient inclusion (T1) and 6-week timepoint (T2) samples as compared to healthy controls. Inclusion versus control samples showed 321428 significant differentially methylated CpGs, and 6-week timepoint samples versus control samples gave 235708 significant differentially methylated CpGs. Hierarchical clustering was performed to visualize global correlation among T1, T2, and healthy control samples. Two major clusters obtained from the analysis showed a cluster with healthy controls and a cluster with patients, which indicated a general separation between COVID-19 patients and healthy controls **(Figure 2A)**. Principal component analysis, an unsupervised learning method, to show variation in the data sets revealed a clear separation of hospitalized COVID-19 patients from healthy controls, revealing a highest variation in data set (PC1) i.e., 32.2% and second most variation (PC2) of 8.8% **(Figure 2B)**. There were exceptions with two patient samples at inclusion (P02, and P03) and three samples at the 6-week time point (P07, P09, and P24), which clustered with the healthy controls, as well as one healthy control sample (HC01) that clustered with COVID-19 patients. The reason for patients clustering with healthy controls or overlap of some patient samples with healthy controls could be the underlying conditions that had led to moderate/severe COVID-19 that was not linked to altered airway/lung pathology or medications that protected the airway compared to the other COVID-19 patients. The latter appears to be the case for all patient outliers besides one since they had been on anti-inflammatory medication for chronic pulmonary conditions. Regarding the healthy control individuals that clustered with the patient samples, we could not find an obvious explanation from the clinical data/history.

**Figure 2:**
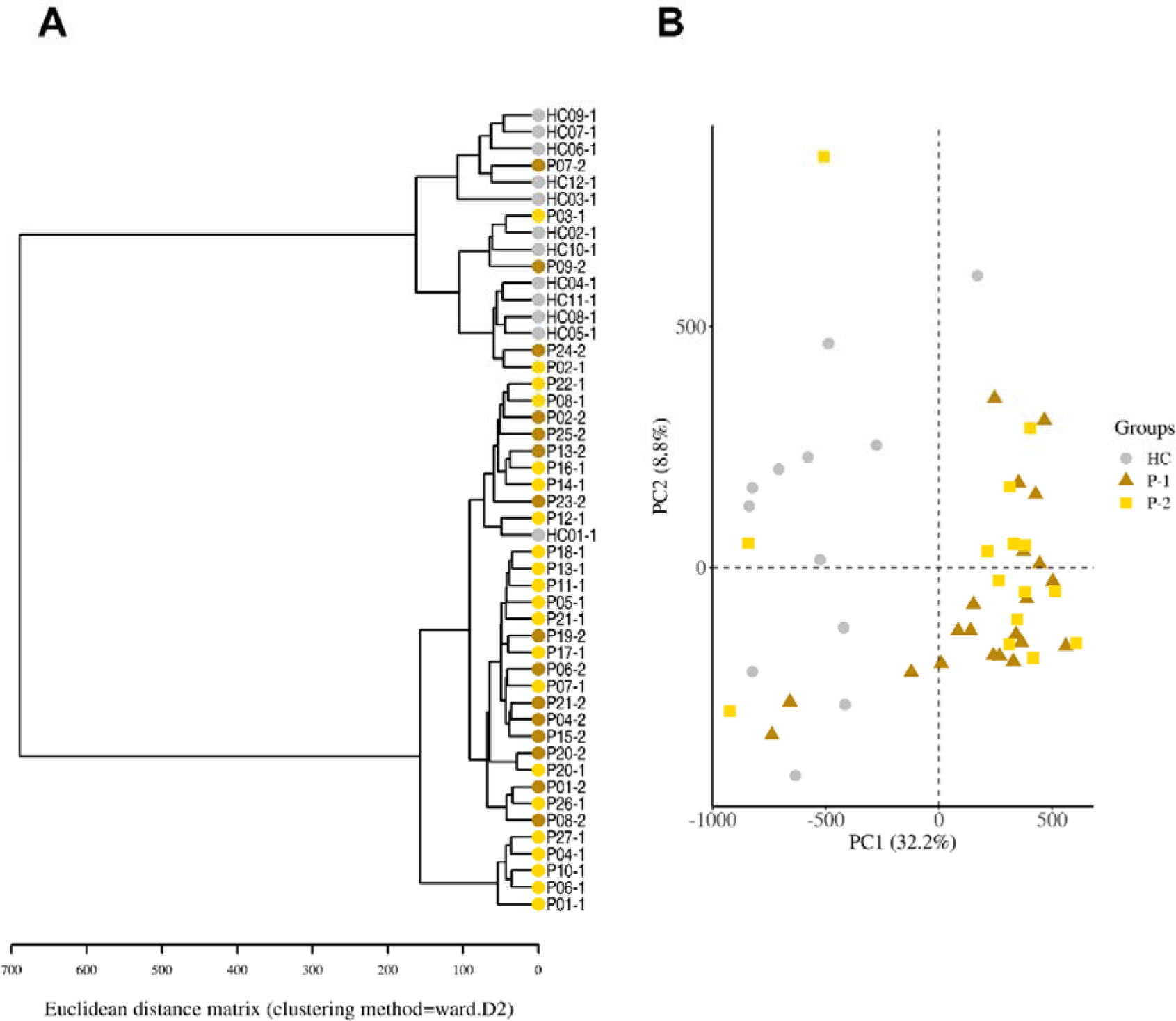
Distinct separation of COVID-19 patients compared to healthy controls. Nasopharyngeal samples (N=36) were collected from 27 COVID-19 patients (P) at inclusion (T1) N=21 and 6-weeks post-inclusion (T2) N=15, and from healthy controls (N=12) (HC) at inclusion of the study. **(A)** Hierarchical clustering using the Euclidean distance performed on the β-value data matrix to visualize global correlation was performed and **(B)** Principal component analysis of normalized β-values of COVID-19 patients and healthy controls.

### Majority of the COVID-19 altered methylated CpGs remained stable over 6 weeks, but a substantial fraction was found only at inclusion

Next, we compared the difference and overlap in CpGs between inclusion versus healthy controls (T1-HC), and between 6-week timepoint versus healthy controls (T2-HC). A unique set of 94382 differentially methylated CpGs were found in the T1-HC that did not exist in the T2-HC, and a unique set of 8609 differentially methylated CpGs in the T2-HC that was not found in the T1-HC **(Figure 3A)**, indicating that most of the differentially methylated CpGs still persisted more than 6-week post infection. This suggests a major difference in DNA methylation patterns between COVID-19 patients and healthy individuals, and that most of the alterations in CpG methylation remained for at least 6-week post infection **(Figure 2** and **3A)**. Based on these findings, we used the pooled inclusion and 6-week timepoint samples versus control samples for additional analysis.

**Figure 3:**
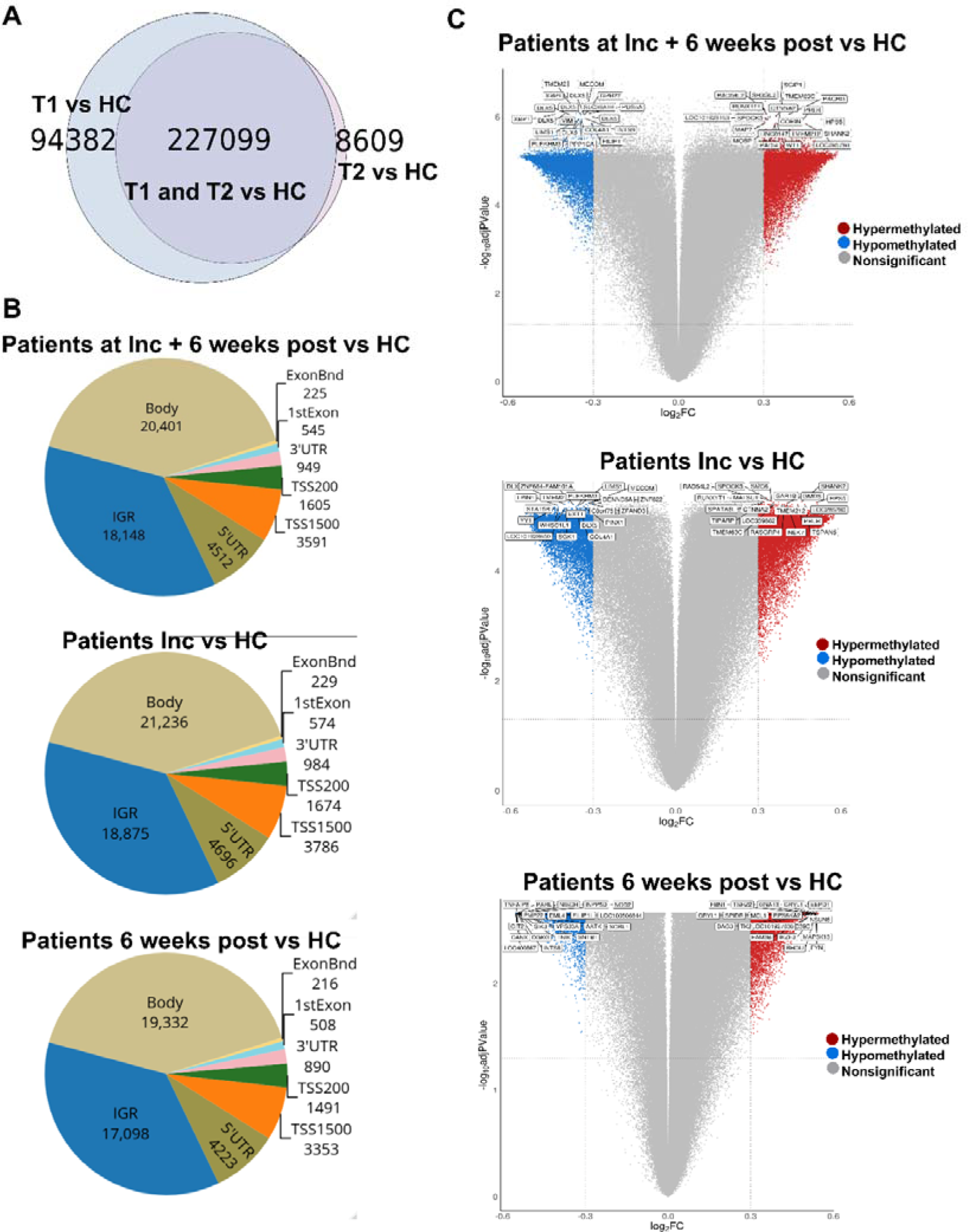
Distribution of differentially methylated CpGs and genes in COVID-19 patients compared to controls. Nasopharyngeal samples (N=36) collected from 27 COVID-19 patients at inclusion (T1) N=21 and 6-weeks post-inclusion (T2) N=15, were assessed for differential methylation within the different genomic regions: Body, Intergenic regions (IGR), 3’ UTR and 5’ UTR, 1^st^ Exon, Exon boundaries (ExonBnd), and the transcriptions start sites TSS200 and TSS1500, and compared to healthy controls (N=12) (HC). **(A)** Venn diagram of unique and common differently methylated CpGs. **(B)** Genomic region distribution for Patients at T1 & T2 vs HC, Patients at T1 vs HC, and Patients at T2 vs HC. Significant set using the Bonferroni-Hochberg (BH)-corrected p-value (p-value BH) < 0.05 and log2FC cut-off (log2FC) set at ±0.3. **(C)** Volcano plots analyzing the log_2_ fold change (log_2_FC) with a cut-off > 0.3 for hypermethylated CpGs and cut-off < 0.3 for hypomethylated CpGs and the use of BH corrected p values < 0.05 cut-off for hypermethylated and hypomethylated CpGs. Top 20 DMGs were annotated using the highest BH corrected p-values highest for hypermethylated and hypomethylated CpGs.

Next, we identified the amount of significant differentially methylated CpGs in the various genomic regions, i.e., gene body (body), intergenic region (IGR), exon boundaries (ExonBnd), 1^st^ Exon, 3’UTR, 5’UTR, and the transcription start sites (TSS) 1500 and 200, in COVID-19 patients versus HCs **(Figure 3B)**. The greatest number of differentially methylated CpGs in the different genomic regions appeared in the body region **(Figure 3B)**. There were no major differences in the levels of the significant differentially methylated CpGs in the different genomic regions between groups, although in general, the 6-week post samples had less significantly differently methylated CpGs in the different regions compared to the inclusion timepoint **(Figure 3B)**.

### Differently hypomethylated and hypermethylated sites/genes with highest significance diverge in the COVID-19 patients between the inclusion and 6-week time points

Methylation can affect gene transcription either by enhancing, decreasing, or silencing transcription ^19, 20^. Hypomethylated and hypermethylated CpG sites specific for COVID-19 were identified based on log_2_FC values with a set cut-off ±0.3 and *p* value_BH_ <0.05 and the top 20 differentially hypomethylated- and hypermethylated CpG sites/genes were emphasized **(Figure 3C**, **Table 2)**. The hypermethylated genes with the lowest -log_10_adjusted *p* value included, SGIP1, SH3GL3, RAD54L2, TMEM63C, and PACRG, associated with the differentially methylated CpGs for hospitalized COVID-19 patients at inclusion and at 6-weeks post-inclusion versus HCs, while the top five hypomethylated genes were, TMEM2, MECOM, XBP1, DLX5, and GRP77 **(Figure 3C, T1 & T2 vs HC)**. The hypermethylated genes with the lowest -log_10_ adjusted *p* value at inclusion (T1) encompassed SPOCK3, SMC6, SHANK2, RAD54L2, and RUNX1T1, and top hypomethylated genes, DLX5, TMEM2, ZNF664-FAM101A, PLEKHM3, MECOM (**Figure 3C)**. By 6-weeks post inclusion (T2), COVID-19 hypermethylation of genes with lowest -log_10_ adjusted *p* value included FBN1, TSHZ2, GNA13, CRYL1, and EEPD1, and hypomethylated genes included TNFAIP8, PARL, NISCH, INPP5D, and NOD2 **(Figure 3C)**. There were no overlapping methylated genes when comparing genes with the highest *p* value_BH_ between inclusion and 6-weeks post-inclusion **(Figure 3C, T1 vs T2)** in COVID-19 patients versus healthy controls. Of note, there was a remarkable decline in the hypomethylated and hypermethylated genes’ -log_10_ adjusted *p* values between the inclusion and 6-weeks post-inclusion, with a decline from ∼5 to 3, indicating that the altered CpG methylation induced by COVID-19 was getting restored to the levels found in healthy, i.e., as before the infection. When exploring the top hypermethylated genes according to the highest fold change in the patients at inclusion compared to healthy controls we found BAG3, EHF, FAM178B, TMX2-CTNND1, and MCC, and for top hypomethylated genes we found NISCH, PMP22, SFMBT2, FAM124A, and FCER1G **(Supplementary Table 1)**. By 6 weeks post-inclusion, COVID-19 hypermethylation of genes with highest fold-change compared to healthy controls included RALGAPA2, PDGFD, C1orf228, MCC and FAM178B, while the top five hypomethylated genes were LOC400867, TNFAIP8, GIT2, CANX and PARL **(Supplementary Table 1)**.

**Table 2:**
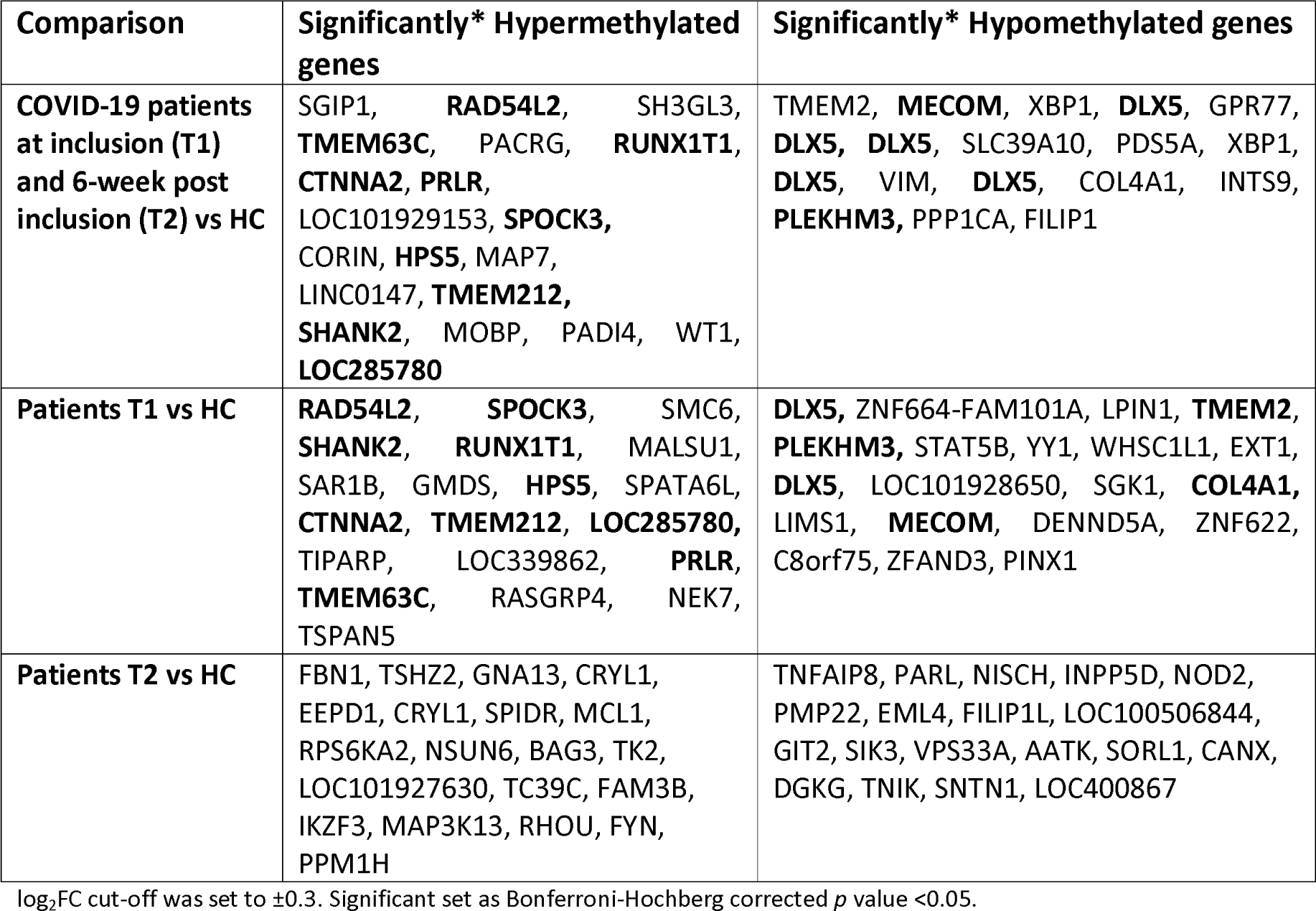
Top hyper- and hypomethylated genes with unique differentially methylated CpGs in COVID-19 patients compared to controls.

### Different methylated CpGs patterns in COVID-19 patients versus healthy controls

For a snapshot of the overall pattern of differently methylated CpGs covering all the different genomic regions in hospitalized COVID-19 patient at T1 and T2 versus HCs, a heatmap with hierarchical clustering of the DNA methylation beta-values was constructed with top 20000 methylated CpGs. This showed a clear difference in methylation between the COVID-19 patients and controls **(Figure 4)**. Following this, we focused on the specific methylation patterns for the transcription start sites TSS1500 and TSS200, as methylations in these areas are good predictor of activation versus silencing of genes ^41^. The DNA methylation beta-value was analyzed using hierarchical clustering with the top 1000 differentially methylated CpGs and the distribution of other associated parameters, timepoint, age and biological sex; and clinical parameters associated with disease, i.e., LDH and C-reactive protein (CRP) **(Figure 5)**. There was a distinct pattern of differently methylated CpGs in the TSS1500 and TSS200, but we found no specific relation to age or biological sex, CRP, or LDH, with the pattern of differently methylated CpGs **(Figure 5)**. The gene body methylation had similar patterns as seen for transcription start sites **(Supplementary Figure 2).** Of note, the COVID-19 patients had, within the selected transcription start sites TSS1500, TSS200, and the gene body, more genes/CpGs with high beta values than the healthy controls, i.e., higher level of methylated CpG sites.

**Figure 4:**
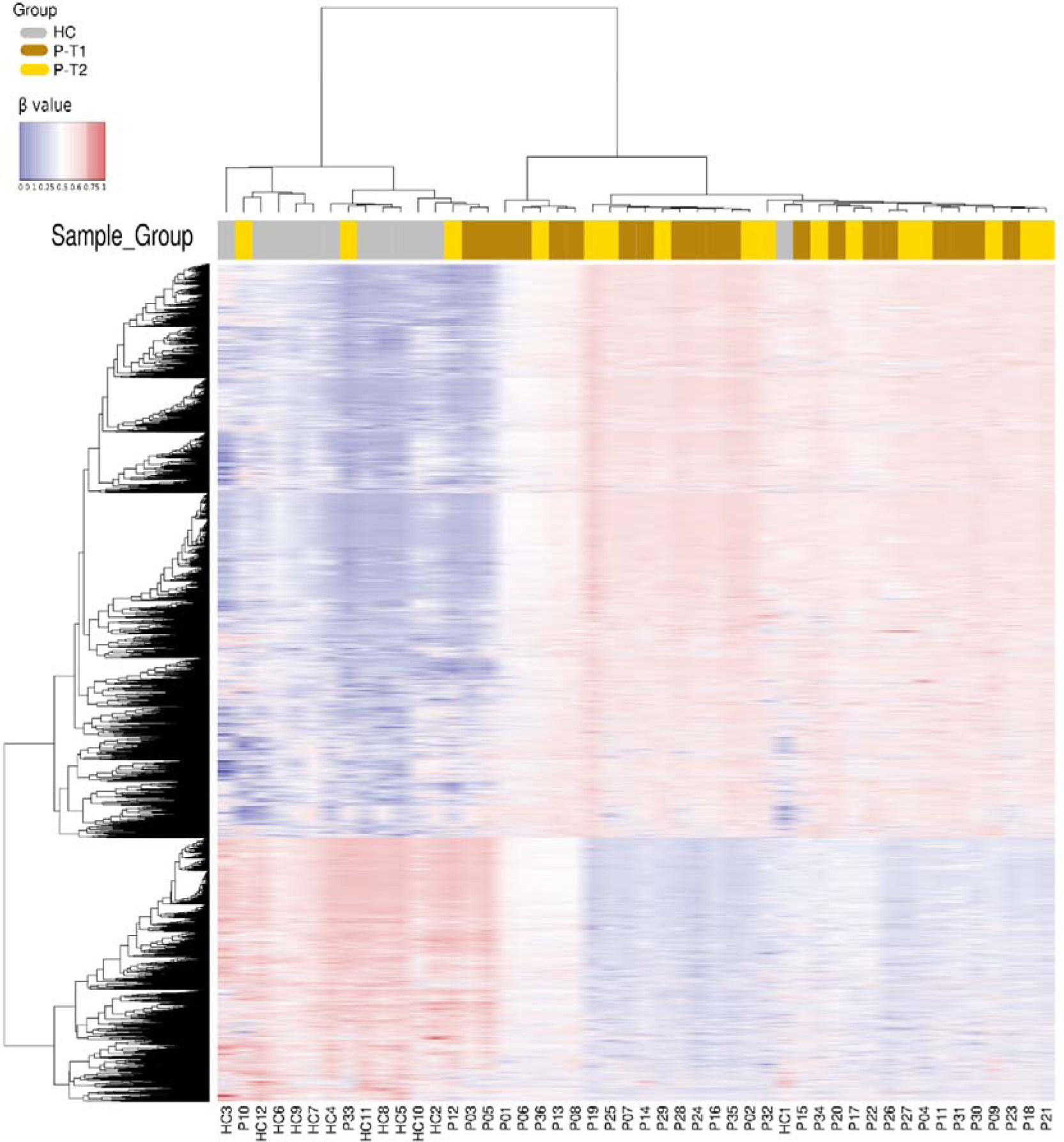
Clear distinctions in the methylation patterns of COVID-19 patients compared to healthy controls. Nasopharyngeal samples (N=36) collected from 27 COVID-19 patients (P) at inclusion (1) N=21 and 6-weeks post-inclusion (2) N=15, and healthy controls (N=12) (HC) were assessed for differential methylation pattern showing the top 20 000 data points hypo and hypermethylated CpGs.

**Figure 5:**
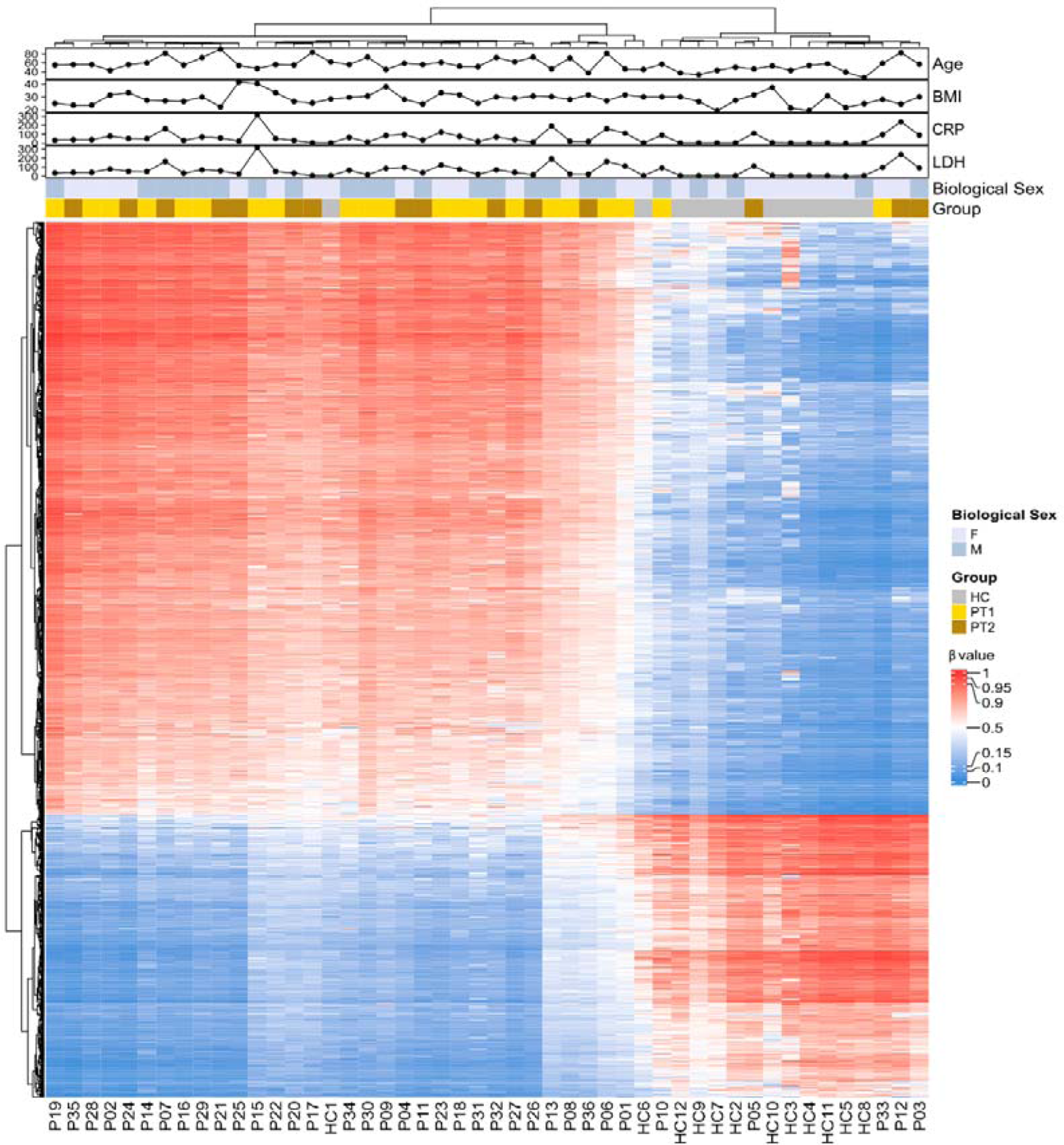
Differentially methylated CpGs between COVID-19 patients and controls in transcription start sites TSS200 and TSS1500. Nasopharyngeal samples (N=36) collected from 27 COVID-19 patients (P) at inclusion (T1) N=21 and 6-weeks post-inclusion (T2) N=15 and healthy controls (N=12) (HC) were assessed in the TSS200 and TSS1500 regions for top 1000 hypo and hypermethylated CpG and 40 differently methylated CPG sites annotated randomly. Hierarchical clustering against sample groups was performed. Top annotation showed the distribution of Sample Groups, Gender, BMI, Age, CRP, LDH.

### Enrichment analysis of genes methylated in the transcription start sites in COVID-19 patients indicated an effect on genes involved in inflammatory and immune responses

To investigate the biological significance of methylated genes in COVID-19 patients, we pooled the top 500 hypomethylated and 500 hypermethylated transcription start site regions TSS1500 and TSS200 encoding genes and performed an enrichment analysis using Metascape database (http://metascape.org) ^42^. The top enriched ontology terms included regulation of cell activation, leukocyte activation, and inflammatory responses **(Figure 6A),** and top biological processes that included multicellular organismal processes, immune system process, and response to stimulus **(Figure 6B)**. The large number of unique differentially methylated genes in the TSSs between the inclusion and 6-week post timepoint (T1 & T2) **(Figure 3A, Supplementary Table 2)**, showed that the initial effects of the viral infection and immune responses occurring early during COVID-19 (T1) did not persist at the later stage (T2). This finding indicates that some of the CpG methylations in the TSSs induced by COVID-19 were short-term. Further enrichment analysis of the unique genes affected at T1 was performed, using the top 50 differently methylated CpGs at the TSSs, according to adjusted *p* value, and fold-change. For T1 versus HC the top-level gene ontology biological processes were Response to stimulus and Cellular process (adjusted *p* value), whereas the top-level gene ontology biological processes according to fold-change differences were responsible for stimulus and growth **(Figure 8D)**.

**Figure 6:**
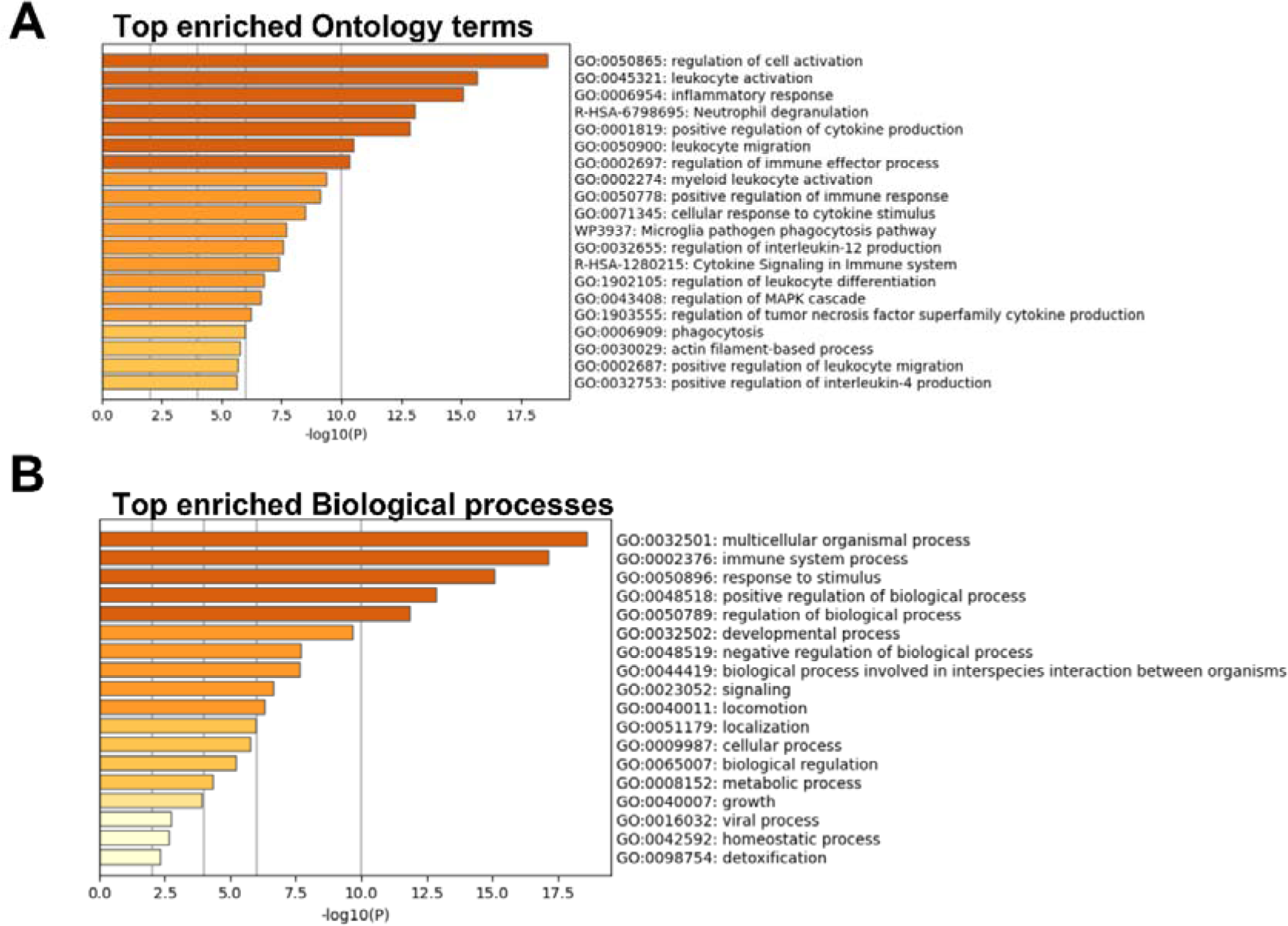
Enrichment analysis for COVID-19-specific differentially methylated genes in transcription start sites TSS200 and TSS1500. Nasopharyngeal samples (N=36) collected from 27 COVID-19 patients (P) at inclusion (1) N=21 and 6-weeks post-inclusion (2) N=15, and from healthy controls (N=12) (HC), were processed and evaluated for DNA methylation. Analysis was performed with Metascape ^42^ for **(A)** enriched terms and **(B)** biological processes for the top 1000 hypo- and hypermethylated genes.

Enrichment analysis in DisGeNET also highlighted delayed myelination, which is commonly caused by viral infections ^43^. Whilst looking at the enrichment according to fold-changes, we found Immune system process and Localization to be the top-level gene ontology biological processes, and an enrichment for infective cystitis, which has been reported to occur in some COVID-19 patients^46^. Further enrichment analysis in TRRUST^44^ indicated the top predicted biological processes for T2 versus HCs, as Multicellular organismal process and Negative regulation of biological process (adjusted *p* value), and an enrichment of the regulation by the tumor suppressor gene TP53. Of note, there was an enrichment for the regulation by IRF-1, highlighting the key role of interferons in COVID-19 ^47^. The two genes of interest that were enriched included at T1 IL-17A, a key cytokine involved in inflammation and healing ^48^, and NUP93 at T2, which is associated with antiviral innate immunity ^49^.

### RNAse1 and RNAse2 were identified as top regulators, and IL-18 was involved in several biological processes in COVID-19 patients

To identify activated and inhibited canonical pathways, and to identify top regulator networks based on all significant hypomethylated and hypermethylated transcription start site regions encoding genes, we used Ingenuity Pathway Analysis (IPA) **(Figure 7)**. The canonical pathways predicted to be activated included PI3K/AKT Signaling, Antioxidant Action of Vitamin C, Th2 Pathway, Natural Killer Cell Signaling, and Neuroinflammation Signaling Pathway. The pathways predicted to be inhibited included S100 Family Signaling Pathway, Leukocyte Extravasation Signaling, Phospholipases, and ERK/MAPK Signaling **(Figure 7A)**. RNAse1 and RNAse2, hydrolytic enzymes catalyzing degradation of RNA, were identified as top regulators in the regulator network with the highest consistency score **(Figure 7B)**. RNAse1 and RNAse2 regulate factors such as pro-inflammatory and anti-inflammatory cytokines leading to the homing and chemotaxis of several immune cells and cell death. Next, we explored another top regulator network, with four regulators, i.e., DNA methyltransferase 3 alpha (DNMT3A), an enzyme important for methylation; Ubiquitin specific peptidase (USP22), a ubiquitin-specific processing protease; the enzyme Caspase 4 (CASP4), autophagy-pyroptosis-related function; and Colony Stimulating Factor 3 (CFS3), cytokine controls the production, differentiation, and function of granulocytes. These regulators altogether contribute to increased interactions with multiple enzymes, and transmembrane proteins, which have an effect on functions relating to degranulation of phagocytes and myeloid cells **(Figure 7C)**. A summary of the major biological pathways and factors demonstrating the role of IL-18, an important pro-inflammatory Th1 polarizing cytokine linked to various processes including chemotaxis and migration of leukocytes, including neutrophils, and degranulation **(Figure 7D)**. IL-18 plays a major role in various infectious, metabolic, and inflammatory diseases by inducing IFN-γ thereby promoting Th1 cell activation and enhancing the cytotoxic activity of CD8+ T cells and natural killer (NK) cells ^50^.

**Figure 7:**
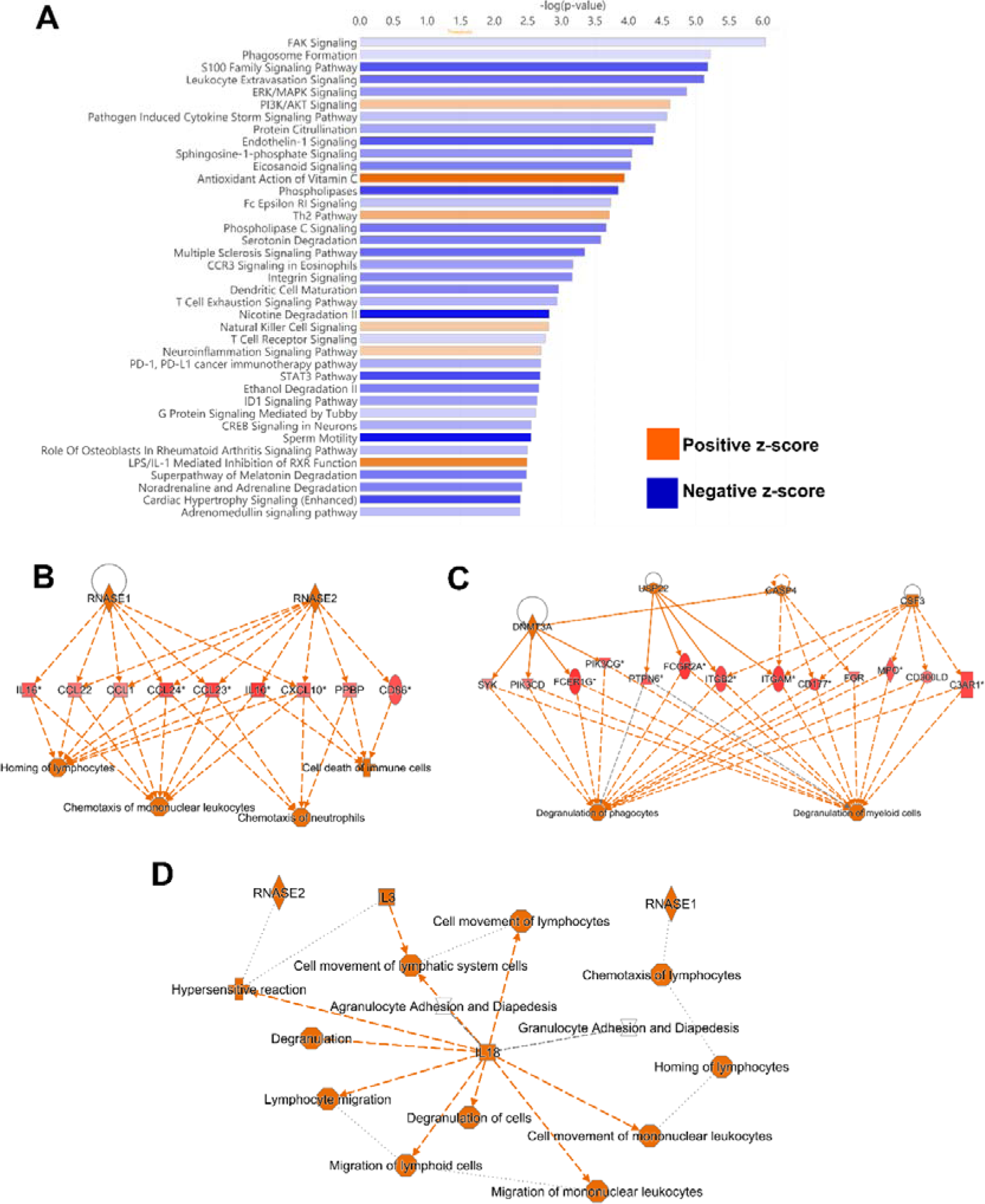
Top canonical pathway and top regulatory networks in hospitalized COVID-19 patients. DNA methylation analysis was performed on nasopharyngeal samples from 27 COVID-19 patients (P) at inclusion (1) N=21 and 6-weeks post-inclusion (2) N=15, and from healthy controls (N=12) (HC). **(A)** Canonical pathway analysis was performed with Ingenuity Pathway Analysis to identify top activated and inhibited pathways linked to the methylated genes located in the transcription start sites 1500 and 200, with a z-score cut-off set to 1. Positive z-score indicated in orange, while negative z-scored indicated in blue. Significant TSS differentially methylated CpGs with a cut-off of +/- 0.3-fold-change was assessed in Ingenuity Pathway Analysis for **(B-C)** top regulator effect networks, and for **(D)** graphical summary.

### Enrichment analysis match of genes methylated in the transcription start sites in COVID-19 patients versus healthy controls compared to other COVID-19 datasets

To examine the similarities with other COVID-19 datasets, we performed an enrichment analysis match of the top 1000 affected genes, consisting of pooled top 500 hypomethylated and 500 hypermethylated transcription start sites regions encoding genes, found in our dataset **(Figure 8)**. Several COVID-19 datasets in the CORONASCAPE database ^42^ were predicted to match, and five datasets were selected based on gene overlap and adjusted logP values. All datasets were from transcriptomic studies on blood and airway samples from SARS-CoV-2 infected patients and in vitro SARS-CoV-2 infected cell line such asprimary human airway epithelial and hypotriploid alveolar basal epithelial cells. There was an overlap in genes between all datasets and many genes were found in several of the datasets **(Figure 8A)**. Of note, there were many genes unique to our data set **(Figure 8A)**. Our DNA methylation data shared enrichment of six genes, i.e., OAS1, CXCR5, APP, CCL20, CNR2, and C3AR1 with the other datasets **(Figure 8B),** of which several have been defined in previous COVID-19 studies ^51, 52, 53, 54^. The enriched top terms defined by the analysis match included regulation of cell activation, leukocyte activation, inflammatory response, regulation of immune effector processes, and neutrophil degranulation **(Figure 8C)**. Furthermore, the top-level biological processes included multicellular organismal processes, immune system process, response to stimuli, and regulation of biological processes **(Figure 8D)**. Taken together, our findings clearly indicate a multilayered activation of responses by SARS-CoV-2 infection.

**Figure 8:**
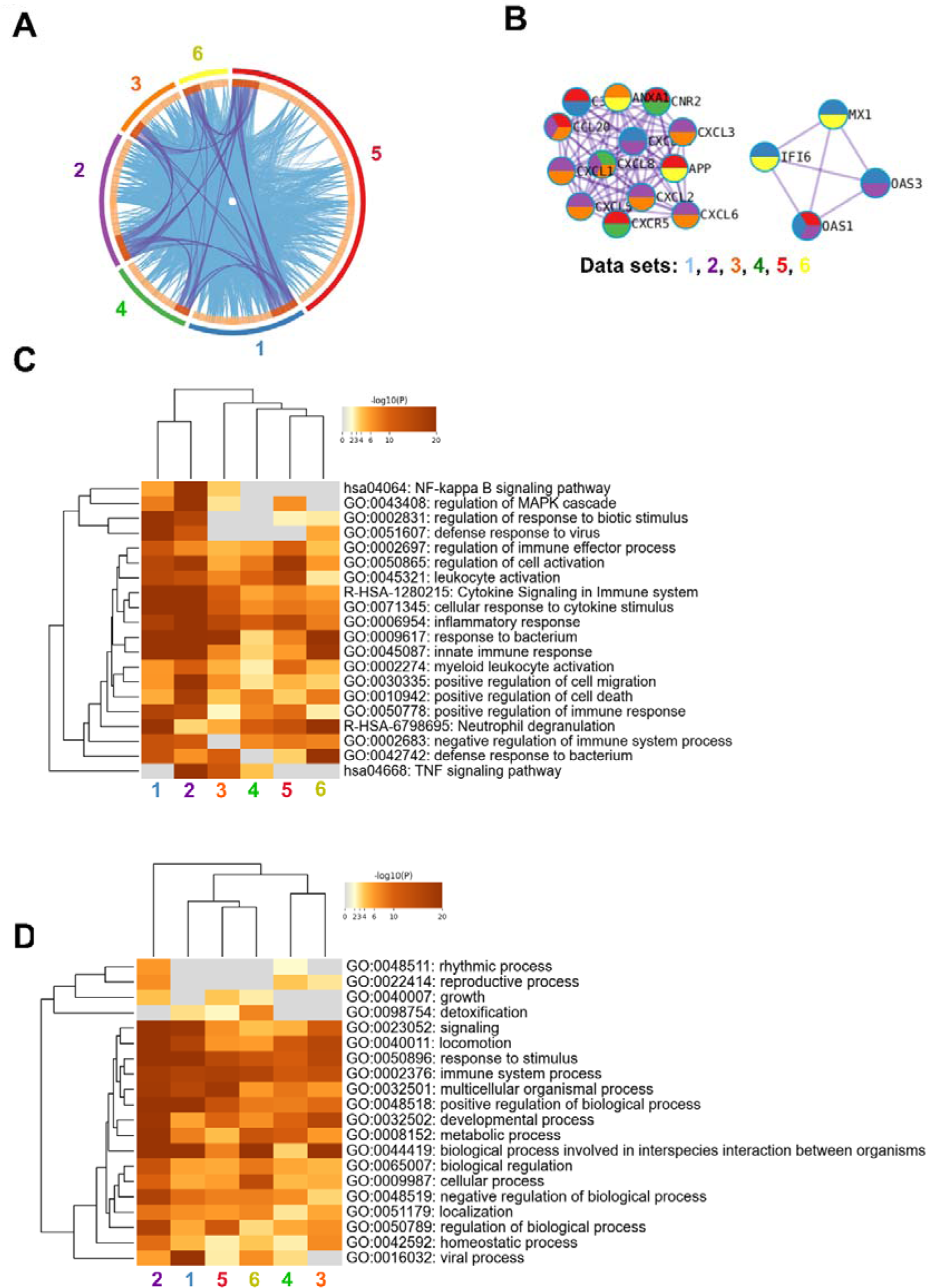
Enrichment match analysis for top 1000 differently methylated genes across COVID-19-specific studies. DNA methylation analysis was performed on nasopharyngeal samples from 27 COVID-19 patients (P) at inclusion (1) N=21 and 6-weeks post-inclusion (2) N=15, and from healthy controls (N=12) (HC). Analysis match was performed against the top 5 selected datasets based on logP values and overlapping genes from COVID-19 studies to assess similarities between enrichment of genes. **(A)** Circos plot for overlapping gene lists and shared term level, where blue curves link genes belonging to the same enriched ontology term. The inner circle represents gene lists, where hits are arranged along the arc. Genes that hit multiple lists are colored in dark orange, and genes unique to a list are shown in light orange. **(B)** Network nodes identifying neighborhoods where proteins are densely connected. Heatmap of **(C)** enriched terms and **(D)** top-level gene ontology biological processes across input gene lists. Each data set has been numbered and color-coded: 1, RNA_Blanco-Melo_Lung_Down; 2, RNA_Sun_Calu-3_0h_Up; 3, RNA Vanderheiden-pHAE_48h_Up; 4, RNA_Lieberman_Nasopharynx_High_vs_Low_Down; 5, gene list from the present study; and 6, RNA_Wilk_B-cells_patient-C6_Up. Enrichment analysis was performed with Metascape and Coronascape ^42^.

## Discussion

Severe infections and conditions will alter the DNA methylation pattern in cells and tissues. Here were nasopharyngeal samples from patients hospitalized with COVID-19 used to identify SARS-CoV-2-induced epigenetic signatures by profiling the DNA methylation patterns at inclusion/hospitalization as well as 6-weeks post-inclusion. We utilized upper respiratory airway samples to explore the effect on the airway as there is evidence that epigenetic alterations that occur in one location or cell type cannot directly be extrapolated for another cell/tissue ^55^. There was a clear separation in the DNA methylation pattern in the airway samples between COVID-19 patients and healthy controls. The high impact exerted by COVID-19 on the DNA methylation pattern in the airway, is in line with other studies that explored this in blood samples from COVID-19 patients ^15, 24, 25, 26, 27, 56^. Interestingly, patients who had undergone anti-inflammatory treatment such as oral/nasal inhalation glucocorticoids for previous/chronic airway manifestations had a different DNA methylation pattern and clustered with healthy controls in the PCA. This suggests that the initial inflammatory response played a major part of the imprinted DNA methylation pattern in the airways and that this did not occur to the same degree among individuals on anti-inflammatory treatment due to less inflammation in the airway. Another explanation could be that the anti-inflammatory drugs alter the DNA methylating effects exerted by SARS-CoV-2 on the airway and the local infiltration of immune cells.

We did not see a clear distinction in DNA methylation between COVID-19 patients with moderate and severe disease when exploring hierarchical clustering. Separate DNA methylation patterns have been demonstrated between mild and severe COVID-19 ^57, 58^.We found some evidence reflecting a restoration of the epigenetic profile in the patients, but even if certain of the COVID-19-induced hypomethylated and hypermethylated CpGs/genes returned to levels found in healthy, there was still major lasting effects on the DNA methylation pattern in the COVID-19 patients. One explanation for the changes in the airway methylation patterns could be altered cell compositions, such as infiltration of immune cells, proliferation of tissue resident cells and damaged epithelia at the initial phases or during active infection versus a more healing phase a few weeks after COVID-19 ^59, 60^.

It’s evident that some of the differently methylated genes that were present only at inclusion in the airway samples, but not at the 6-week post inclusion time point, such as OAS1 and OAS3 were part of the innate type 1 interferon responses to viruses. These genes have been confirmed to be affected in COVID-19 patients and are linked to clinical outcome ^10, 25^. In blood samples, others have shown that some DNA methylation alterations are present one year after SARS-CoV-2 infection in hospitalized patients ^61^. Genome areas with genes and CpGs that returned to normal levels were connected to viral responses such as type I IFN signaling, whereas areas associated with cell activation, leukocyte activation, lymphocyte activation, immune system remained altered over time ^61^. Our findings are in line with these previous findings in blood^61^ with regards to DNA methylation returning to normal levels in genes involved in viral responses such as the OAS1, whereas genes involved in e.g. leukocyte activation, and immune system remained altered.

We saw alterations in DNA methylation in all genomic regions in the COVID-19 patients, but focused on the deeper analysis of the DNA hypermethylation and hypomethylation patterns in the transcription promotor regions as this more easily can be translated to silenced and activated genes, respectively ^62^. Further, a previous study found that differential patterns of COVID-19 DNA methylation in blood occurred primarily in the promoter regions of immune-related genes ^27^.

The top enriched ontology and biological processes predicted by the pathway analysis in the COVID-19 data set included regulation of cell activation, leukocyte activation, inflammatory responses, leukocyte migration, immune system process, and response to stimuli, which is in accordance with previous COVID-19 studies ^25, 63^. Signaling pathways predicted to be affected included the S100 signaling pathway and PI3K/AKT Signaling. S100 family genes have been identified as prognostic markers of severe COVID-19 ^64^ and the PI3K/Akt intracellular pathway might be an important signaling pathway in the cytokine storm induced by SARS-CoV-2 and also in COVID-19 coagulopathies ^65^.

The endogenous antimicrobial polypeptides RNAse1 and RNAse2 were defined as top regulators in the COVID-19 DNA methylation data set. Zechendorf et al reported that elevated RNASE1 levels were linked to renal dysfunction among ICU-admitted COVID-19 cases ^66^. Others found increased RNAse2 levels in COVID-19 patients with critical disease ^67^. The role(s) RNAses may play in SARS-CoV-2 infection is largely unknown. RNASEs might play a role regulating the innate immune response activated by SARS-CoV2 by recognizing and degrading single- and double-stranded RNA molecules and thereby limiting the TLR3 and TLR7/8 ligands, i.e., extracellular viral RNA, released by Infected dying cells ^68^

An analysis of the major biological pathways and factors demonstrated a key role for IL-18, an important proinflammatory cytokine induced by IFN-γ and linked to various processes including chemotaxis and migration of leukocytes, inflammasome activation, pyroptosis and cytotoxic activity of CD8+ T cells and NK cells ^69^. IL-18 has been shown to play a major role in various infectious, metabolic, and inflammatory disorders ^50^. COVID-19 with elevated levels of IL-18 has been associated with increased severity and mortality ^70, 71^. Furthermore, IL-18 SNPs/mutations, such as IL18-105G>A, have been shown to be protective against severe COVID-19 ^72^.

Whilst exploring some of the other genes affected by COVID-19, we found that the IL-10 gene was hypomethylated in COVID-19 patients at inclusion/hospitalization, indicating an active IL-10 response early on during infection and it has been shown to play a part in COVID-19 pathogenesis, both during acute infection and post/long COVID-19 ^73, 74^. Genes involved in innate immune responses such as NOD2, KLK7 and genes involved immune regulation, such as SEMA4D, INPP5D, TOX, IKZF3, were identified among the differently methylated genes with the highest fold change in COVID-19 patients. The epigenetic imprinting by the SARS-CoV-2 infection on these genes involved in innate responses and immune regulation is important determinates for the disease development ^75^.

Furthermore, we observed alterations of TSSs for genes involved in protein transcription, translation, folding, quality control, post transcriptional modification affected e.g. CANX, GPN3, RBM47, METTL21A, and BAG3 and in neurological development/disorders e.g. PARL, ADK, TSHZ2, ASAM10, and AATK. Infections and inflammation impact all steps in the production, control, and modification of new proteins and SARS-CoV2 has a direct impact on protein translation, and protein trafficking ^76^, and need post translational modifications ^77^. Regarding neurological effects of COVID-19 there are reports that have demonstrated direct effects on the nervous system and connection between long COVID-19 ^78, 79^ and exploring our data set’s matches with other data sets revealed mutual enrichment of six genes, i.e., OAS1, CXCR5, APP, CCL20, CNR2, and C3AR1 ^10, 51, 52, 53, 54, 80, 81, 82, 83, 84^. The enriched top terms defined by the analysis match included regulation of cell activation, leukocyte activation, inflammatory response, regulation of immune effector processes, and neutrophil degranulation. The genes and the biological processes/responses defined in the enrichment analysis are commonly found in numerous COVID-19 studies {Blanco-Melo, 2020 #113{Wilk, 2020 #117}{Vanderheiden, 2020 #119} and reflect the effects the viral infection and subsequent immune response exert on the host.

Severe conditions such as sepsis are connected to changes in DNA methylation patterns in white blood cells in areas with genes involved in, e.g., inflammatory pathways, innate and adaptive immune response, which are reflected in altered gene expression ^85^. Subsequently, a relevant comparison of our data would be with other severe infectious diseases to determine the DNA methylation alteration that is specific for COVID-19. In addition, a larger cohort could also increase the ability to determine with higher precision unique DNA methylations and patterns for COVID-19 and it would be of interest to verify some findings with functional tests.

In conclusion, we found a clear distinct DNA methylation pattern in the COVID-19 patients compared to healthy controls and that most but not all altered methylations lasted for more than 6 weeks post inclusion. in the COVID-19 patients compared to controls. There was an enrichment of genes with altered methylation at transcription start sites associated with Inflammation and immune system in the COVID-19 data set, which might reflect an attempt to dampen the inflammation and immune reaction.

## Supporting information

Govender et al Supplementary tables and figures

## Author contributions

MG, FH, CS, handling of study samples. ML and MG designed the experiments. MG conducted experiments. MG, ML, JD, SN, YKY, JK, EMS and ML analyzed the data. AN-A, AJH, MH, JS, JN, SN, and ML were involved in establishing the cohort study. MG, JD, EMS, SN, VV, RS, and ML were involved in the writing of the initial manuscript draft. MG, EMS, JD, JK, SN, SR, VV, FH, CS, JN, YKY, AN-A, AH, MH, JS, and ML were involved finalizing the manuscript for submission. All authors read and approved the final manuscript.

## Acknowledgements

We are grateful for all patients and healthy donors who participated in this cohort study. We thank all individuals involved in the study, including coordinators, and all healthcare personnel from the Clinic of Infectious Diseases, and the Intensive Care Unit at the Vrinnevi Hospital, Norrköping, Sweden. We would like to thank Patrick Müller and David Brodin at the Bioinformatics and Expression Analysis Core facility, Karolinska Institute. We would like to acknowledge the Bioinformatics Core Facility, Faculty of Medicine and Health Sciences and Clinical Genomics Linköping, SciLife Laboratory, Department of Biomedical and Clinical Sciences, Linköping University, for assistance with bioinformatic analyses.

Our sincere thanks to Annette Gustafsson for helping with the study coordination and sample collection. We would like to thank Mario Alberto Cano Fiestas and Robin Göransson for their assistance with sample processing.

## Ethical statement

The studies involving human participants were reviewed and approved by Swedish Ethical Review Authority. The patients/participants provided their written informed consent to participate in this study.

