## Supplementary material for "Altered DNA methylation pattern contributes to differential epigenetic immune signaling in the upper respiratory airway of COVID-19 patients": Govender et al Supplementary tables and figures

**Supplementary Table 1: Top 30 differentially hypomethylated- and hypermethylated CpG sites/genes according to fold change**

| | COVID-19 Patient T1 vs HC | | COVID-19 Patient T2 vs HC | |
|---|---|---|---|---|
| | **Hypermethylated** | **Hypomethylated** | **Hypermethylated** | **Hypomethylated** |
| **1** | **BAG3** | **NISCH** | RALGAPA2 | **LOC400867** |
| **2** | EHF | **PMP22** | PDGFD | TNFAIP8 |
| **3** | **FAM178B** | **SFMBT2** | C1orf228 | **GIT2** |
| **4** | TMX2-CTNND1 | FAM124A | **MCC** | **CANX** |
| **5** | **MCC** | **FCER1G** | **FAM178B** | **PARL** |
| **6** | **FYN** | **LOC100506844** | **COX10** | **NISCH** |
| **7** | **MCL1** | **CANX** | SLC25A25 | **DGKG** |
| **8** | FBXO32 | **SIK3** | TACC2 | **INPP5D** |
| **9** | MIR21 | GPN3 | **CLIC6** | **NOD2** |
| **10** | **CRYL1** | C10orf12 | **TMCC1** | INTS8 |
| **11** | CUX1 | LOC101927865 | **CRYL1** | **SIK3** |
| **12** | **TMCC1** | ADK | **SPIDR** | TNIK |
| **13** | **NSUN6** | **LOC400867** | **MCL1** | SNTB1 |
| **14** | LINC01589 | C12orf79 | RPS6KA2 | VPS33A |
| **15** | **RHOU** | RBM47 | **BAG3** | AATK |
| **16** | **TK2** | **DGKG** | **TK2** | **PMP22** |
| **17** | ADAM10 | **NOD2** | **LOC101927630** | SORL1 |
| **18** | **TTC39C** | HEXA | FAM3B | EML4 |
| **19** | **CRYL1** | **PARL** | FBN1 | FILIP1L |
| **20** | **GNA13** | C14orf34 | **TSHZ2** | **LOC100506844** |
| **21** | **TSHZ2** | SEMA4D | **TTC39C** | MBTD1 |
| **22** | **MAP3K13** | **RAPGEF2** | **GNA13** | PDGFRB |
| **23** | **SPIDR** | METTL21A | **CRYL1** | LOC285626 |
| **24** | **LOC101927630** | **INPP5D** | **EEPD1** | **SFMBT2** |
| **25** | TOX | EIF3H | **IKZF3** | LOC100506895 |
| **26** | **PPM1H** | **GIT2** | **RHOU** | **FCER1G** |
| **27** | **IKZF3** | BLM | **FYN** | CD44 |
| **28** | **EEPD1** | IL10 | **PPM1H** | **RAPGEF2** |
| **29** | **CLIC6** | KLF7 | **MAP3K13** | MAML3 |
| **30** | **COX10** | TCF7L2 | **NSUN6** | C11orf42 |

Fold change = FC, Healthy control = HC, T1 = inclusion/hospitalization time point, T2 = 6 week post inclusion

**Supplementary Table 2: Top 50 unique differently methylated genes at T1 and T2 vs HC**

| T1 vs HC (adjPv) | T1 vs HC (FC) | T2 vs HC (adjPv) | T2 vs HC (FC) |
|---|---|---|---|
| ARHGEF7 | LRRC2 | CDH4 | JRK |
| NR0B2 | FABP7 | LOC100506142 | ATHL1 |
| IL17A | COG5 | ZFAT | CDK2 |
| ADAMTS7 | LHX6 | H1FOO | TMEM85 |
| MIR3607 | ENOX1 | C6orf106 | NUP93 |
| LOC101929260 | GLYATL3 | C5orf13 | SMAP1 |
| LIMS2 | FUT8 | EBF3 | THRSP |
| GATA4 | C20orf78 | BCL3 | TRIM22 |
| ABCB10 | ERMN | RFX8 | RELB |
| SEPT9 | DPYS | C8orf88 | C11orf75 |
| OR5D18 | TRIM44 | C3orf22 | RWDD3 |
| SMPD3 | C3orf26 | AGER | HOXA5 |
| OPCML | GRIP2 | LOC100507250 | C13orf15 |
| MIR1261 | MTPN | FAM24B | GLOD4 |
| FAM69A | SLC15A5 | LTBR | GLUL |
| POLRMT | LOC102724053 | TMEM170A | PCDHB15 |
| TSPAN3 | TMEM139 | FAM189A2 | TMEM85 |
| SLC6A19 | MRPL15 | EHF | ELF5 |
| TTLL6 | MIR1-2 | INSL3 | PSKH2 |
| ARL5B | EFHA1 | RAB3IP | PIWIL1 |
| ATE1-AS1 | C11orf52 | RNU6-52P | PCDH18 |
| TTLL3 | ALS2CR12 | GPR132 | C10orf88 |
| ZNF211 | FABP7 | KIAA1549 | ZNF84 |
| WFIKKN2 | SLC25A25 | SLC39A13 | LINC00442 |
| GRXCR1 | COG5 | GPNMB | IL17A |
| RSPH6A | TTC3 | OR52M1 | SLC13A3 |
| NARS | FUT8 | XPO1 | PCDHB6 |
| ARPP21 | FAM69A | ADGRG1 | LOC100505920 |
| METTL5 | OR2A5 | MNS1 | IFIT3 |
| EFS | ZNF385D | LRRTM1 | PLBD1 |
| ANKRD39 | ZNF385D | CMTM3 | RNF4 |
| ZNF341 | TBCE | C9orf23 | NKAIN1 |
| RECQL | KIF2B | SCPEP1 | SFXN3 |
| VPRBP | CDH18 | MOG | C16orf63 |
| SFRP1 | LOC105616981 | GPR108 | ODF3B |
| SCN9A | TEK | C5orf24 | GUF1 |
| NEK1 | SPRR3 | NXPH4 | TMEM189-UBE2V1 |
| LINC01504 | CSN1S2BP | CIPC | CHGB |
| FEM1C | HIF1AN | SMYD1 | CHCHD5 |
| PIP5K1A | KLHDC7A | C4orf34 | C1QL3 |
| ARAP2 | PRICKLE1 | C11orf75 | NMNAT2 |
| URI1 | PHACTR1 | KCNIP2 | UCP2 |

| | | | |
|---|---|---|---|
| ZFP30 | DPYS | TBC1D30 | TAF1B |
| NSMCE1 | C13orf26 | PRKCSH | MRPS22 |
| GGA2 | SOD3 | NADK | SH3BP2 |
| ZFAND1 | IDE | ICA1 | CACNA1B |
| MAVS | REG3G | CLASP2 | MED15 |
| PLIN4 | GMNN | NEUROG1 | STAC2 |
| YTHDF2 | SYBU | PDZRN3-AS1 | TCEA1 |
| NCMAP | GPR21 | ZNF461 | M1AP |

Fold change = FC, Healthy control = HC, T1 = inclusion/hospitalization time point,
T2 = 6 week post inclusion, adjPv= adjusted P value

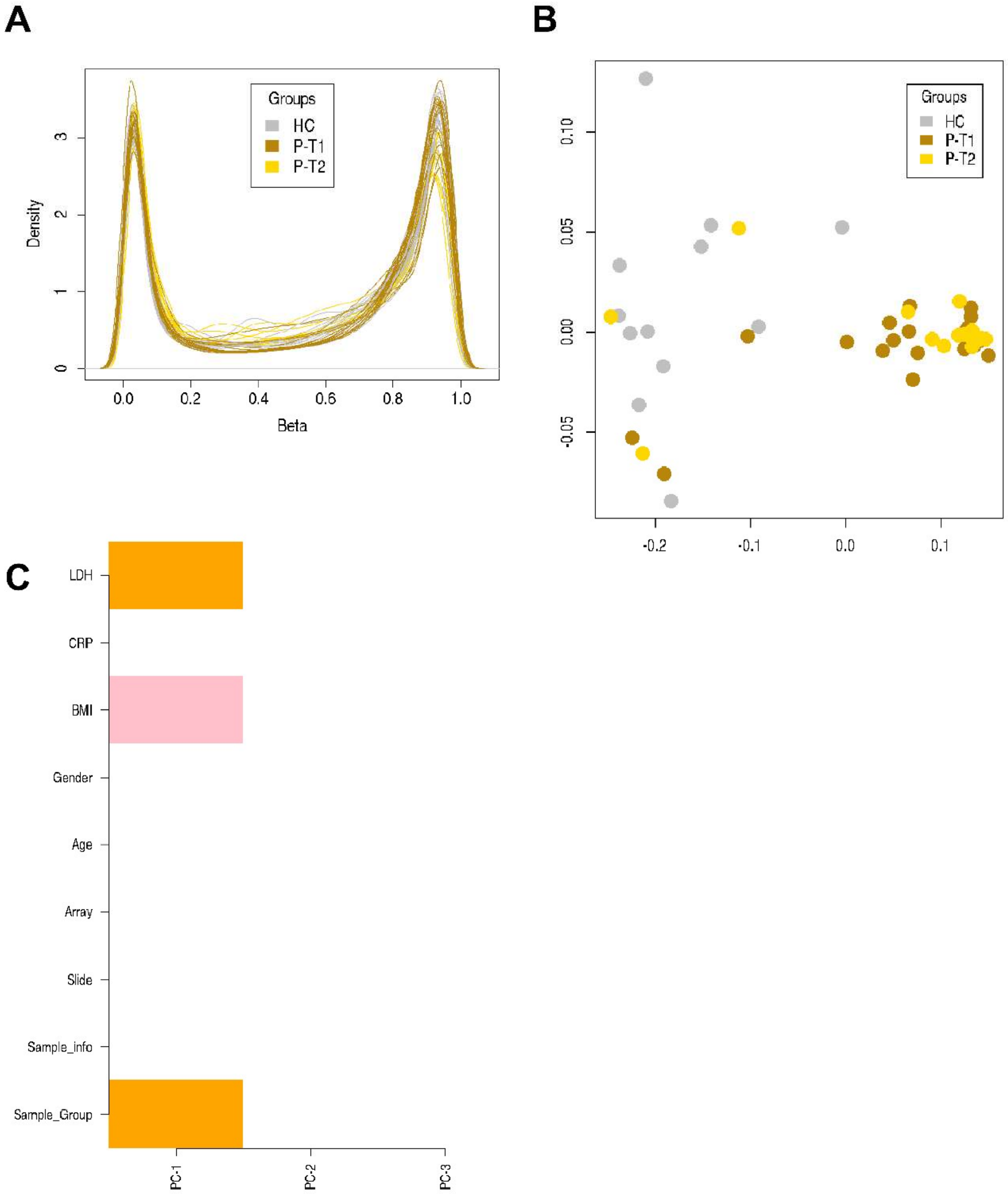


**Supplementary Figure 1. Distinct separation of COVID-19 patients compared to healthy controls.**

Nasopharyngeal samples (N=36) were collected from 27 COVID-19 patients (P) at inclusion (T1) N=21 and 6-weeks post-inclusion (T2) N=15, and from healthy controls (N=12) (HC) at inclusion of the study. **(A)** Initial quality control assessment with beta distribution**, (B)** multidimensional scaling plot for T1, T2 and HC samples, and **(C)** Components of variation/confounding factors in the methylation dataset. The effect of these factors was adjusted by deconvolution and LDH was disregarded due to being a patient-related factor, and the dataset was adjusted for the BMI.

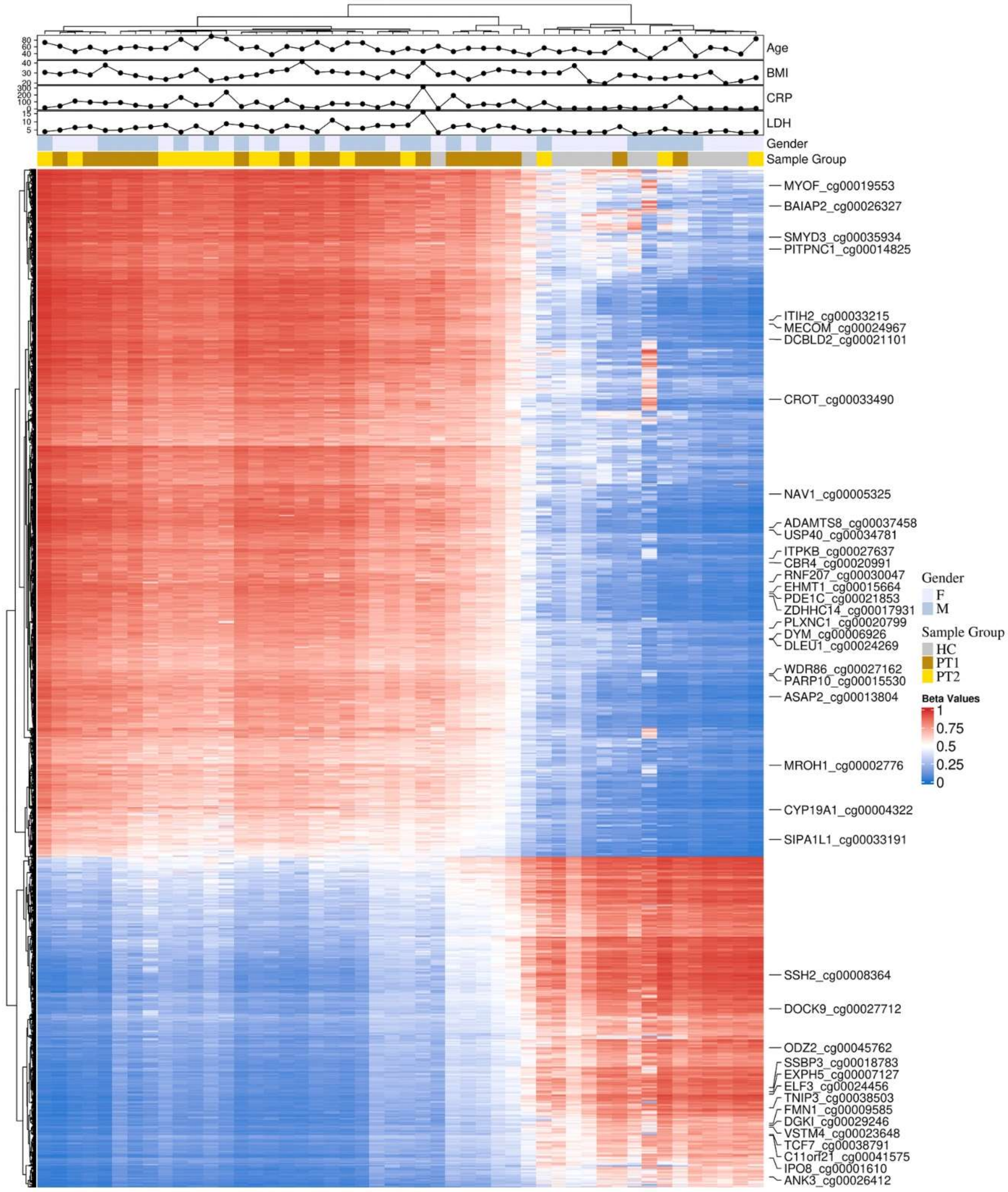


**Supplementary Figure 2. Differentially methylated CPGs between COVID-19 patients and healthy controls in gene body region.**

Nasopharyngeal samples (N=36) collected from 27 COVID-19 patients (P) at inclusion (T1) N=21 and 6-weeks post-inclusion (T2) N=15 and healthy controls (N=12) (HC) were assessed in the gene body regions for top 1000 hypo and hypermethylated CpG and 40 differently methylated CPG sites annotated randomly. Hierarchical clustering against sample groups was performed. Top annotation showed the distribution of Sample Groups, Gender, BMI, Age, CRP, LDH.
